# Reduction of retinal ganglion cell death in mouse models of familial dysautonomia using AAV-mediated gene therapy and splicing modulators

**DOI:** 10.1101/2023.05.22.541535

**Authors:** Anastasia Schultz, Shun-Yun Cheng, Emily Kirchner, Stephanann Costello, Heini Miettinen, Marta Chaverra, Colin King, Lynn George, Xin Zhao, Jana Narasimhan, Marla Weetall, Susan Slaugenhaupt, Elisabetta Morini, Claudio Punzo, Frances Lefcort

**Affiliations:** Department of Microbiology and Cell Biology, Montana State University, Bozeman, MT; Department of Ophthalmology, Neurobiology & Gene Therapy Center, University of Massachusetts Chan Medical School, Worcester, MA; Center for Genomic Medicine, Massachusetts General Hospital Research Institute, Boston, MA; Department of Neurology, Massachusetts General Hospital Research Institute and Harvard Medical School, Boston, MA; Department of Biological and Physical Science, Montana State University Billings, Billings, MT; PTC Therapeutics, Inc., South Plainfield, NJ 07080

## Abstract

Familial dysautonomia (FD) is a rare neurodevelopmental and neurodegenerative disease caused by a splicing mutation in the Elongator Acetyltransferase Complex Subunit 1 (*ELP1*) gene. The reduction in ELP1 mRNA and protein leads to the death of retinal ganglion cells (RGCs) and visual impairment in all FD patients. Currently, patient symptoms are managed, but there is no treatment for the disease. We sought to test the hypothesis that restoring levels of Elp1 would thwart the death of RGCs in FD. To this end, we tested the effectiveness of two therapeutic strategies for rescuing RGCs. Here we provide proof-of-concept data that gene replacement therapy and small molecule splicing modifiers effectively reduce the death of RGCs in mouse models for FD and provide pre-clinical data foundation for translation to FD patients.

## Introduction

Familial dysautonomia (FD) is a recessive autonomic and sensory neuropathy. Over 99% of FD patients are homozygous for the “founder" mutation in intron 20 of the *ELP1* gene (c.2204 + 6T > C; formerly called “*IKBKAP*”), causing the “skipping” of exon 20 from the mature mRNA coding sequence [1, 2]. This mutant mRNA is then targeted for non-sense mediated decay [3]. The mis-splicing occurs in a tissue-specific manner, with peripheral neurons being most impacted, producing almost no functional protein [4, 5]. The hallmarks of FD include reduced pain and temperature sensation, gait ataxia, cardiovascular instability, swallowing impairment, gastrointestinal dysfunction, and eventual blindness [6]. As patients age, visual impairment becomes one of the most debilitating symptoms, given that it is accompanied by loss of balance and gait ataxia. The optic neuropathy in FD results from the progressive death of retinal ganglion cells (RGCs) and the loss of their axons from the nerve fiber layer (NFL) in the retina’s macular region [7-9].

It is necessary to develop therapeutics to thwart the loss of RGCs occurring in all FD patients. Being a monogenic disease, FD is a strong candidate for genetic therapies. Gene replacement therapies for retinal diseases show tremendous potential due to the accessibility and immune-privileged status of the eye [10]. Adeno-associated viruses (AAVs) are efficient viral vectors in preclinical and clinical studies for delivering functional genes in hereditary retinal diseases [10-12]. In addition to gene therapy, a potent strategy for treating genetic diseases is using splicing modulator compounds (SMCs) that correct mutations at the messenger RNA level. Such a modifier was recently approved to restore correct splicing in the neurological disorder spinal muscle atrophy [13, 14]. SMCs effectively reduce disease phenotypes in FD mouse models [15-17]. However, until now, no gene therapy approaches have been used to introduce wild-type copies of *ELP1* to correct disease phenotypes. Here we present pre-clinical data demonstrating the effectiveness of gene therapy and a novel splicing compound in preventing RGC death in FD mouse models.

## Results

Using two complementary FD mouse models, mice were treated either with (1) an intravitreal injection of an AAV2 (highly tropic for RGCs) engineered to express a wild type copy of *ELP1* [18-20] (Figure 1A) or (2) a novel small molecule splicing modulator to restore correct *ELP1* splicing [16] (Figure 1B).

**Figure 1:**
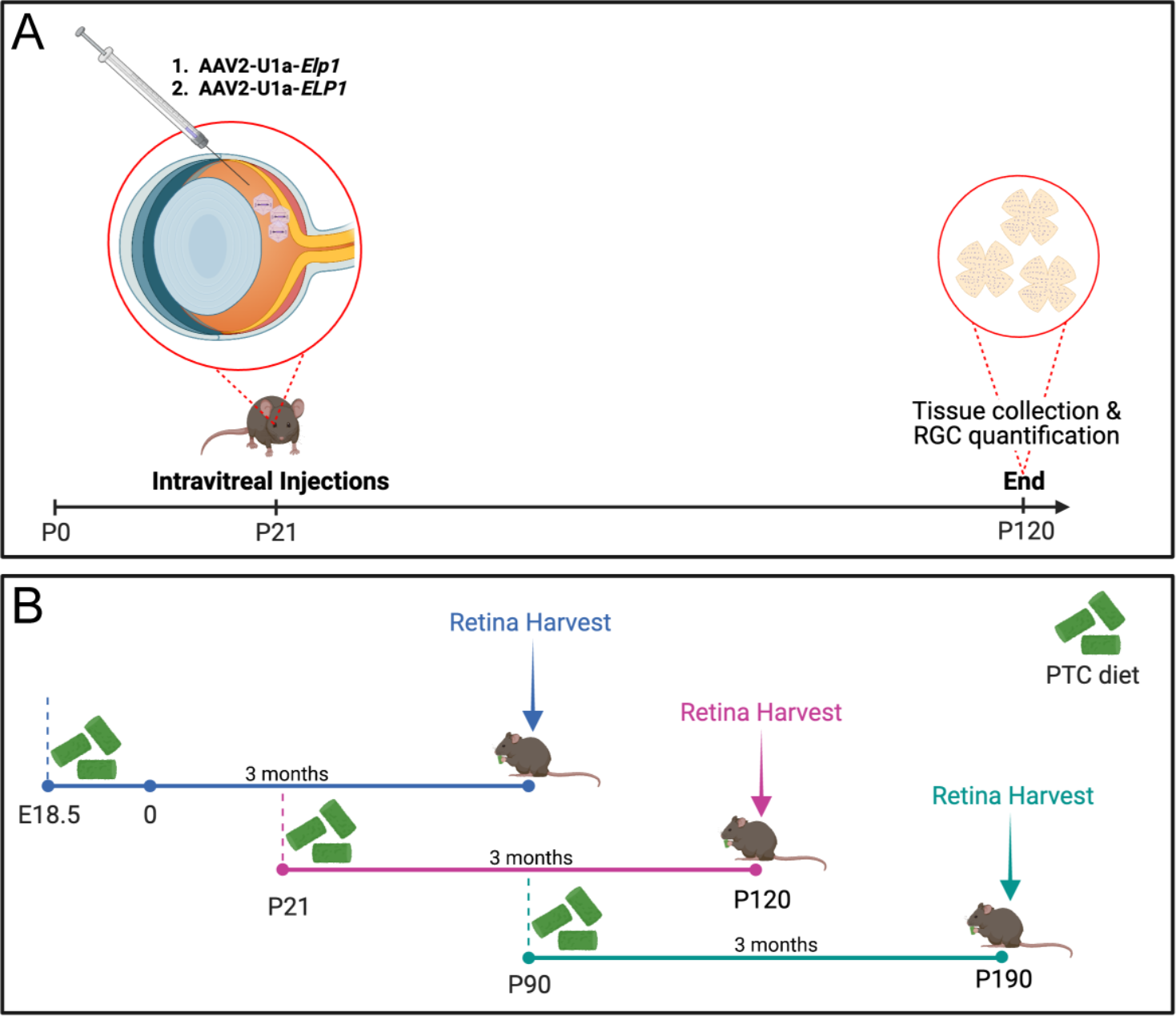
Experimental workflow. (A) Mice were intravitreally injected with an AAV2-vector in both eyes at P21, and the retinal ganglion cell (RGC) number was quantified 3 months later. An experimental or control virus was injected into the vitreous humor of the mouse eye, where it can diffuse into the retina and target RGCs. (B) FD mutant mice were divided randomly into two groups: control and experimental. The control group was fed a standard chow diet, and the experimental group with a specially formulated chow diet containing a novel splicing modulator compound, PTC680. Groups were started on treatment at three different ages: E18.5, P21 or P90. After 3 months on either the control or experimental diet, retinas were analyzed and quantified for exon 20 inclusion and RGC numbers, respectively. Created with BioRender.com

To test whether a gene replacement strategy could prevent the death of RGCs in our previously established *Pax6-cre;Elp1^loxp/loxp^* mouse model [21], we generated two AAV2 vectors, one expressing the murine *Elp1* gene *(mElp1)* and the other expressing eGFP, both driven by the U1a promoter (see Methods). We first determined that the AAV2-U1a-*mElp1* could drive the expression of *Elp1* in the mouse retina and increase Elp1 protein expression *in vitro* (Supplemental Fig. 1).

Next, cohorts of *Pax6cre;Elp1^loxp/loxp^* mice received intravitreal injections of AAV2-U1a-*mElp1* or control (eGFP) vector at postnatal day 21 (P21) before any significant loss of RGCs is observed (Ueki et al., 2018). Three months following injection, RGCs were quantified by flat-mount analysis (Figure 2A). This time point was selected since prior work established that, by this age, 40-60% of RGCs have died. We observed a significant response with a 137% increase in RGC number (Mean: 5696 ± 252, p < 0.0001) with the higher AAV2-U1a-*mElp1* titer (x10^9^ vg/eye) and a 131% increase in RGC number (Mean: 5563 ± 172, p < 0.0001) with the lower AAV2-U1a-*mElp1* titer (x10^8^ vg/eye) in *Pax6cre; Elp1^loxp/loxp^* retinae compared to untreated *Pax6cre; Elp1^floxp/loxp^* retinae (Mean: 2404 ± 405) (Figure 2B). Interestingly, eyes injected with the control AAV2-U1a-*eGFP* showed an 82% increase in RGCs (Mean: 4386 ± 534) compared to the untreated *Pax6cre; Elp1^loxp/loxp^* (Figure 2B). However, the *mElp1-virus* significantly increased RGC numbers by 30% (p = 0.04) above the eGFP-injected eyes. This suggests that (1) injection of a virus in a dose-dependent manner alone induces a protective response in the retina, most likely by stimulating a wound response, but (2) that the *Elp1* virus increased RGCs significantly above the virus-alone-induced protective response. These data provide promising evidence that gene replacement therapy, with further optimization, could be used to ameliorate RGC death in FD.

**Figure 2:**
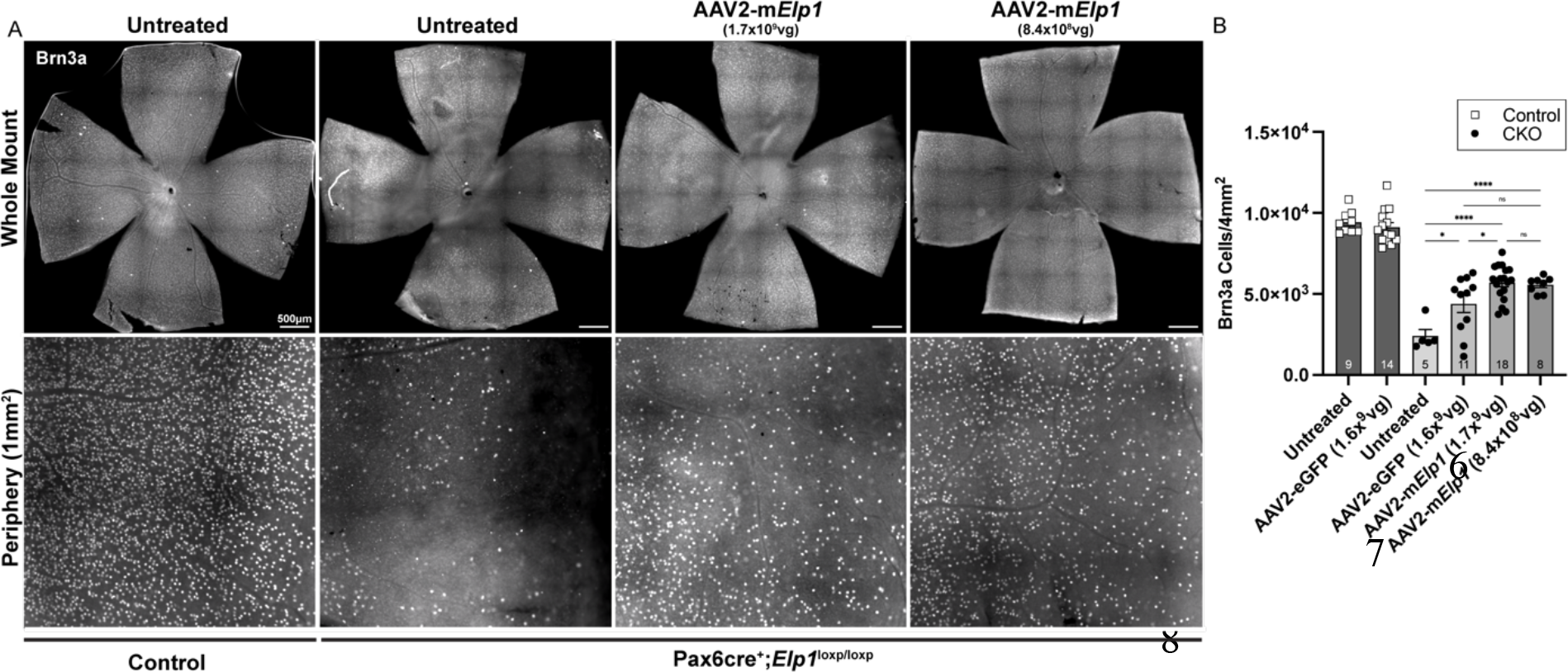
Viral transduction of murine *Elp1* rescues RGCs in a mouse model of FD. (A) Representative retinal whole mount (top) and peripheral retina (bottom) 3 months after intravitreal injection with AAV2-U1a-m*Elp1.* RGCs were immunolabeled using an antibody against Brn3a. Scale bar, 500µm. (B) Quantification of Brn3a + RGCs from control mice (dark gray) compared to FD mice (lighter gray gradient) receiving injections at P21 with either the *eGFP* or experimental *mElp1* virus. Brn3a + cells were counted in a 1mm^2^ area in each quadrant of the peripheral retina. The adjusted p-values are displayed. ns p = 0.2, 0.99, *p = 0.02, 0.04, **** p < 0.0001, one-way ANOVA with Tukey’s multiple comparisons follow-up test. Data are shown as average ±SEM, and each data point represents an individual retina. The sample size for individual groups is represented by *n* within each bar.

Based on these data, we generated an AAV2 vector driving the expression of the human *ELP1* gene (*hELP1)*. Again, we confirmed that the AAV2-U1a-*hELP1* could increase ELP1 protein *in vitro* (Supplemental Fig. 2A,B) and drive *ELP1 mRNA* expression in wild-type mouse retinas *in vivo* (Supplemental Fig. 2C). Subsequently, *Pax6cre;Elp1^loxp/loxp^* and littermate controls were intravitreally injected with three different viral titers of the human *ELP1* virus. All mice received bilateral injections at P21, and RGCs were quantified three months later (Figure 3A). There was a significant increase in RGC survival in the *Pax6cre;Elp1^loxp/loxp^* retinae that received the highest viral titer (5.4×10^8^ vg/eye), showing a 39% increase in RGC number (Mean: 4925 ± 439, p = 0.02) (Figure 3B). There was no significant difference between the uninjected retina and those injected with the AAV2-U1a-*eGFP* construct, which was used at a lower dose than in the mouse Elp1 virus experiments. To visualize the AAV2-U1a-*hELP1* infected RGCs, we double immunolabeled retinae with antibodies to RGCs and human ELP1. In the retinae infected with AAV2-U1a-*hELP1,* approximately 26% of RGCs expressed the human ELP1 protein (Figure 3C), visually confirming that driving *ELP1* expression can efficiently produce human ELP1 protein.

**Figure 3:**
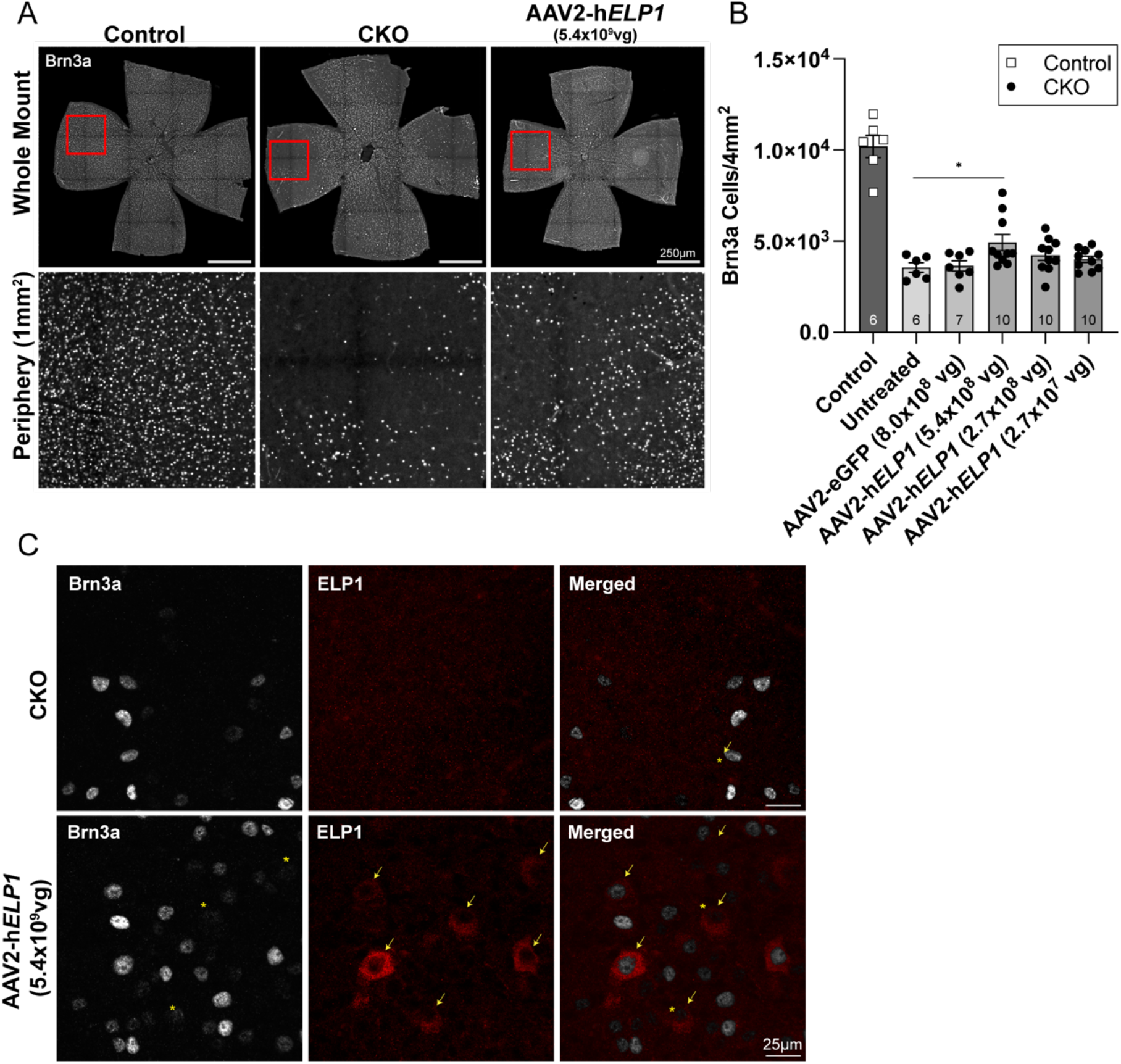
*In vivo* expression of human *ELP1* prevents loss of RGCs in a mouse model of FD. (A) Representative retinal whole mount (top) and peripheral retina (bottom) 3 months after intravitreal injections with *hELP1.* RGCs were immunolabeled using an antibody against Brn3a. Scale bar, 250µm. (B) Quantification of Brn3a + RGCs from control littermate mice (dark gray) compared to FD mice (lighter gray gradient) receiving injections at P21 with either the eGFP or experimental *ELP1* virus. Brn3a + cells were counted in a 1mm^2^ area in each quadrant of the peripheral retina. (C) Representative images of Brn3a+ RGCs (gray, yellow asterisks represent lower Brn3a intensities) and the human ELP1 protein expression (red, yellow arrows) 3 months after intravitreal injections into mutant mice. Scale bar, 25µm. The adjusted p-value is displayed. *p = 0.02, unpaired Mann-Whitney U test. Data are shown as average ±SEM, and each data point represents an individual retina. The sample size for individual groups is represented by *n* within each bar.

Another validated approach for treating genetic disease is using orally administered small molecules that modify splicing [22, 23]. To test the efficacy of a novel small splicing compound PTC680, an analog to the highly specific *ELP1* splicing modulator PTC258 [16], we generated a new hybrid mouse model in which we introduced the human *FD ELP1* original founder splice mutation (*TgFD9)* transgene into the *Pax6cre;Elp1^loxp/loxp^* conditional knockout mouse model. This new model recapitulates the FD retinal phenotype while recreating the tissue-specific mis-splicing observed in FD patients. To assess the therapeutic effect of PTC680, we administered an oral treatment formulated to dose each *TgFD9*;*Pax6cre;Elp1^loxp/loxp^*mouse 1.2 mg/kg/day. Treatment started at three different time points, (a) Embryonic day 18.5 (E18.5), (b) P21, and (c) P90, to test the therapeutic efficacy throughout the lifespan and disease progression (Figure 1B). In all cases, mice consumed either vehicle or PTC680 special chow for 3 months. RGCs were quantified at the end of each 3-month treatment regimen using retinal flat-mount analysis (Figure 4A). PTC680 treatment increased RGC number by up to 57% depending on the age at which treatment began (Figure 4B). The most significant increase in RGCs occurred when treatment began at the youngest age (E18.5). Notably, a significant rescue of RGCs occurred even when treatment started at 3 months old. Thus, daily consumption of PTC680 prevents significant RGC death in both mature and immature retinas.

**Figure 4:**
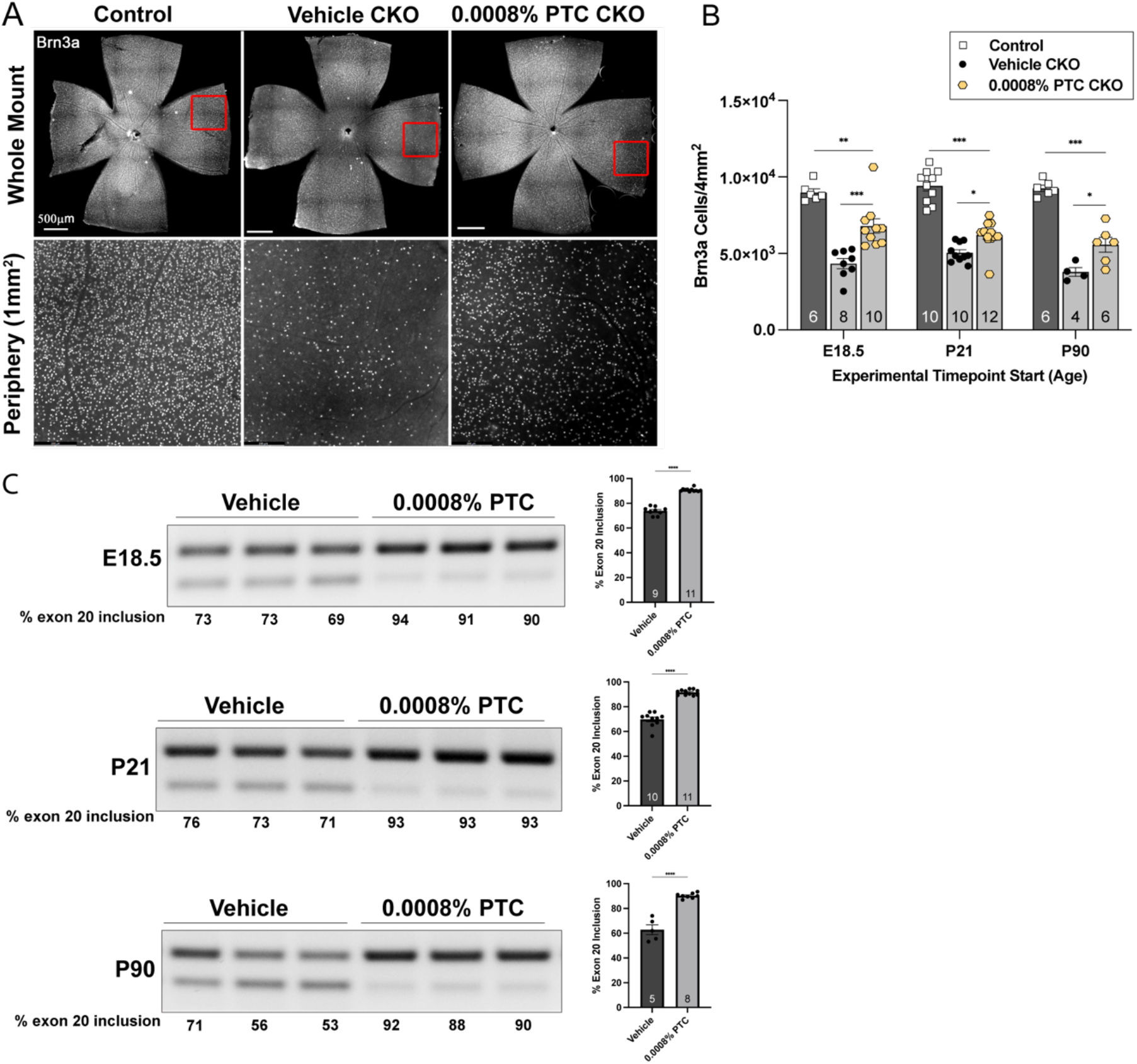
Daily PTC680 consumption protects retinal ganglion cells from cell death and increases full-length *ELP1* transcript in a new retinal FD mouse model. (A) Representative retinal whole mount (top) and peripheral retina (bottom) after daily consumption of PTC680 for 3 months. Scale bar, 500µm. (B) Quantification of Brn3a + cells from control littermate mice (dark gray) and FD mice (light gray) on the vehicle or experimental chow. All mice consumed chow for 3 months, but each experiment started when the mice were at a different age: E18.5 (**p = 0.004, ***p < 0.001), P21 (*p = 0.01, ***p = 0.0000) or P90 (*p = 0.02, *** = 0.0000), to test the efficacy of starting treatment at different time points within disease progression. The adjusted p-values are displayed, one-way ANOVA with Tukey’s multiple comparisons follow-up test (C) Representative splicing analysis of human *ELP1* transcripts and percent of exon 20 inclusion in retinae from vehicle-treated (dark gray) and PTC 0.0008%-treated (light gray) FD mice at the different experimental start times. The adjusted p-values are displayed. ****p < 0.0001, two-tailed unpaired Student’s *t*-test. Data are shown as average ±SEM, and each data point represents an individual retina. The sample size for individual groups is represented by *n* within each bar. This experiment was conducted three times.

To assess whether RGC rescue was associated with increased splicing efficiency of the mutant FD *ELP1* gene, we analyzed *ELP1* splicing in retinal tissue from vehicle- and PTC680-treated FD mice. As demonstrated with PTC258 [16], PTC680 significantly increases *ELP1* exon 20 inclusion in the FD retina (Figure 4C), providing evidence that this novel splicing modulator mitigates a primary FD phenotype by increasing *ELP1* in the retina.

## Discussion

We demonstrate that both an intravitreally-delivered RGC-specific gene transfer and the systemic delivery of a novel small molecule splicing modulator can prevent the progressive death of RGCs. This is the first gene therapy study in an FD model and lays the foundation for a translational path for mitigating RGC loss in FD patients. Additionally, we show that in a new mouse model, the oral administration of the novel PTC680 compound significantly improved RGC survival throughout multiple stages of disease progression. These data are encouraging as most FD patients do not report vision loss until they are teenagers.

Our data show that approximately 30% of RGCs were infected with the human *ELP1* virus and expressed the human ELP1 protein, which is in the same range as the percentage of RGCs rescued (39%). Moreover, we provide the first evidence that morphology is not disrupted in RGCs expressing virally transduced *ELP1*, suggesting that overexpression of human *ELP1* is not harmful to retinal neurons.

These data justify the further optimization of an *ELP1*-vector for translation to a clinical setting. Potential paths forward include optimizing the AAV2 capsid properties via chemical modifications or assembling mosaic capsids consisting of different serotypes could improve targeting specificity and tropism to achieve higher transduction efficiency [24-26]. Alternatively, concatamerization of dual AAV vectors [27-31] can be considered as the *ELP1* transgene is large (4 kb), making packaging challenging. This would provide more flexibility in promoter choice to maximize transduction efficiency in RGCs.

FD is a complex developmental and degenerative disease, yet because it is caused by a single splicing-associated point mutation, it is an excellent candidate disease for both gene therapy and small molecule splicing modulators. In summary, the data presented here provide a rationale for translating gene replacement therapy and splicing modulator compounds into the clinic to prevent the progressive optic neuropathy that plagues all FD patients.

## Materials and Methods

### Animals

All mice were housed in the Animal Resource Center at Montana State University and all animal use protocols were approved by the Montana State University Institutional Animal Care and Use Committee. The generation of the *Pax6cre^+^;Elp1^flox/flox^*CKO is described in Ueki et al., 2018. Briefly, retina-specific *Elp1* CKO mice were generated by crossing Elp1 floxed (International Knockout Mouse Consortium, Wellcome Sanger Institute, UK) and αPax6 promoter-driven Cre (Pax6-Cre) mice [32], which were a gift from the original founders (Drs Peter Gruss and Ruth Ashery-Padan). This is a useful mouse model to study the FD optic neuropathy due to the restricted deletion of *Elp1* only in retinal neurons with a time course of RGC loss consistent with patients while maintaining a phenotypically healthy mouse.

A new hybrid mouse model, the *TgFD9*;*Pax6cre;Elp1^loxp/loxp^* was generated to analyze the splicing compound in a mouse model that expressed the FD founder mutation on a mouse background that was null for mouse *Elp1*. To this end, we crossed our original *Pax6-cre;Elp1^loxp/loxp^* mice, with *FD9/FD9; Elp1^loxp/loxp^* transgenic line that expresses nine copies of the human FD *ELP1* mutation [33]. To ensure that crossing in the *FD9* transgene did not, by itself, rescue the loss of RGCs in the *Pax6cre;Elp1^loxp/loxp^* model, we quantified RGCs in both models at 3 months and found a ∼55% reduction, indicating that the low levels of *ELP1* transcribed from the human gene was not sufficient to prevent the death of RGCs.

The sample size for all experiments was based on a statistical power analysis of 90% with an expectation to confidently detect a 20% difference between experimental and control groups. Both male and female mice were used at the ages indicated (E18.5 up to 6 months), and all controls were littermate *Pax6cre^-^;Elp1^loxp/loxp^.* Mice were randomly assigned to a treatment group based on sex and genotype.

### AAV vector design and production

All AAV viruses were made at the Vector Core in the Gene Therapy Center at the University of Massachusetts Medical School. AAV2 was selected based on its reported tropism for mouse retinal ganglion cells and its use and tolerance in the human eye [34, 35] and preliminary experiments by Yumi Ueki et al (unpublished) revealing its higher expression efficiency in retinal ganglion cells in our mouse line compared to AAV9. Each virus expressed the full-length sequence for either mouse *Elp1* (Origene <u>MC202501</u>, NM 026079) or human *ELP1* (Origene RC2076868), ELP1 (NM_003640), and both were driven by a murine small nuclear RNA promoter (U1a). The U1a promoter was selected based on its small size (∼250 bp) and ability to effectively transduce cells in the central nervous system [36].

### Cell culture

Human embryonic kidney cells (HEK293) were cultured in Dulbecco’s Modified Eagle Medium (DMEM, Gibco #12430-054) supplemented with 10% fetal bovine serum (FBS, Gibco #16000-036), 100 U/ml penicillin/100 µg/ml streptomycin (Corning, #30-002-CI), and 2 mM L-Glutamine (Gibco, #25030-081). HEK293 cells submitted to American Type Culture Collection (ATCC) for STR profiling were an 88% match for the HEK293 cell line, CRL-1573.

### In vitro AAV transduction

HEK293 cells were seeded onto a 12-well plate at a density of 2.5×10^4^ cells/well for 24hr. Two viral solutions were diluted (1) AAV2-U1a-h*ELP1* stock to 2.7×10^9^ VG/ml in DMEM and (2) AAV2-U1a-eGFP stock to 8.8×10^9^ VG/ml in DMEM. The *ELP1* solution was serially diluted to three different multiplicities of infection (m.o.i.) at 20K, 100K, and 400K. M.O.I. was calculated by dividing the initial seeding density by the total viral particles. At 24 hours, the complete medium was removed and supplemented with the viral medium. Cells proliferated for 6 hours in the viral medium. At 30 hours, the viral medium was removed and replaced by the complete medium. Cells were harvested 4 days post-transduction (PTD).

### Western Blot

The western blot analysis was performed as previously described *(Ueki et al.,2018*). Briefly, cells were extracted on ice with lysis buffer (25 mM Tris-HCL, pH 7.5, 150 mM NaCl, 1 mM EDTA, 1% NP40, 5% glycerol) containing 1X HALT protease inhibitor cocktail, 1 mM PMSF, and 10 µM Leupeptine. A Bradford 1X protein-dye reagent (Bio-Rad #5000205) was used to determine the protein concentration. 25 µg of protein was separated by electrophoresis in an SDS-polyacrylamide gel, and the proteins were transferred onto a PVDF membrane. The membrane was blocked for 1 hour in 5% non-fat dry milk (NFDM) in 1X Tris-buffered saline containing 0.1% Tween-20. The membrane was incubated with primary antibodies overnight at 4°C. The primary antibodies to rabbit anti-Elp1 (1:1500) (Anaspec, #AS_54494)) and rabbit anti-GFP (1:2000) (Invitrogen, RRID: AB_221569) were diluted in 1X TBS containing 5% BSA and 0.1% Tween-20. Following primary incubation, membranes were probed with a secondary antibody (Horseradish peroxidase-conjugated goat anti-rabbit IgG (Jackson Laboratories, #111-035-144), 1:10,000 dilution, in 1X TBS containing 5% NFDM and 0.1% Tween-20. Quantitative analysis was performed using Image J (NIH, USA) software, and bands were normalized to total protein and the housekeeping gene *Gapdh* (1:10,000, Millipore, NP_002037).).

### RT-qPCR

Real-time PCR (RT-qPCR) was used to quantify mRNA collected from snap-frozen retinal tissue. Total RNA was isolated and purified using the Direct-zol RNA mini-prep kit (Zymo Research, #R2050). The RNA concentration and purity was measured using BioTek Epoch 2 microplate spectrophotometer. 25 ng of purified RNA was reverse transcribed using SuperScript IV VILO mastermix (Invitrogen, # 11756050). qPCR was performed with cDNA equivalent of 25ng of transcribed RNA using SYBR Green master mix (Bio-Rad, #336501) using the 7500 Fast Real-Time PCR system (Fisher Scientific, Applied Biosystems, #4351106). The cycling condition was performed per the manufacturer’s protocol (Bio-Rad). All primers (Supplemental file 1) were designed using NIH Primer-Blast tool. The relative expression analysis was performed using the delta delta CT method (ΔΔCT) and expressed as fold change normalized to the average of the two housekeeping genes.

### RT-PCR analysis of *ELP1 transcripts*

Retinae were snap-frozen in a dry-ice bath with 95% EtOH upon collection. Tissues were homogenized in ice-cold TRI reagent (Molecular Research Center, Inc., Cincinnati, OH, USA), using a TissueLyser (Qiagen). Total RNA was extracted using the TRI reagent procedure provided by the manufacturer. RNA quality and yield was determined using a Nanodrop ND-1000 spectrophotometer. Reverse transcription was performed with 500ng of total RNA, random primers, and Superscript™III reverse transcriptase (Invitrogen). PCR was performed for the splicing analysis with the 3 µl of cDNA in a total volume of 20 µl with GoTaq Polymerase 2X (Promega) and 32 amplification cycles. We used human-specific *ELP1* primers – forward, 5’ - CCTGAGCAG CAATCATGTG- 3’; reverse, 5’ -TACATGG TCTTCGTGACATC- 3’ to amplify human *ELP1* expressed from the transgene. The PCR products were separated on 1.5% agarose gels for 2.5 hours at 90V. The relative amounts of WT and mutant (Δ20) *ELP1* spliced isoforms in a single PCR were determined using ImageJ and the integrated density value for each band as previously described [37, 38]. The relative proportion of the WT isoform detected in a sample was calculated as a percentage.

### Intravitreal injections

Three-week-old mice were anesthetized by isoflurane inhalation. Prior to anesthesia the pupil was dilated with one drop of phenylephrine (Phenylepherine Hydrochloride Ophthalmic Solution, USP 10%; National Drug Code, [NDC] 42702-103-05, Paragon, BioTeck, Inc) and one drop of tropicamide (Tropicamide Ophthalmic Solution, USP 1%, [NDC] 17478-102-12, Akorn Pharmaceuticals).

Two different intravitreal injection techniques (*Manual Hamilton Syringe* and *WPI-UMP3 with Mirco2T Injector*) were used to determine the optimal system for viral delivery. Each eye was treated as an independent experimental endpoint. All control-treated eyes were injected with an AAV2 preparation expressing eGFP at 1.6×10^9^ or 8.0×10^8^ viral genomes. Experimental-treated eyes were tested in a dose-dependent manner to optimize transduction efficiency and RGC rescue. Experimental-treated eyes receiving the AAV2-U1a-*Elp1* were injected with 1.6×10^9^, 1.77×10^9^, 8.3×10^8^, or 8.6×10^8^ viral genomes of murine *Elp1.* Experimental-treated eyes receiving the AAV2-U1a-*ELP1* were injected with either 5.4×10^8^, 2.7×10^8^, or 2.7×10^7^ vector genomes of human *ELP1*.

#### Manual Hamilton Syringe

Using forceps, the eyelids were pushed back to expose the cornea. With a 32-G, a small hole was made at the margin of the cornea and sclera. This was necessary because the A 34-G needle (#W1690678) had a blunt tip. The 5 µl calibrated Hamilton #65 needle was inserted at the margin of the sclera and cornea and into the vitreous cavity and 1 µl of the viral solution was injected slowly into the cavity. The needle was left in position for another 15s and removed slowly.

#### WPI-UMP3 with Mirco2T Injector (#0916C)

Using forceps, the eyelids were pushed back to expose the cornea. A 36-G beveled needle (NF36BV) attached to a 10 µl Hamilton NANOFIL syringe (WPI, lot# 08C) was inserted at the margin of the sclera and cornea and into the vitreous cavity. Each eye was injected with 600 nL of AAV preparation bilaterally with a 200 nL/sec delivery rate. The needle was left in position for another 10s to allow the pressure in the injection system to equilibrate with the pressure in the eye and removed slowly. After all surgeries, Puralube® Vet Ointment (Dechra Veterinary Products) was placed on the eye per IACUC guidelines. Mice were monitored and kept on a 37°C heating pad until awake and recovered and then transferred to their home cage and monitored for 1 to 2 hours. Mice remained in their home cage with cage mates until analysis, 3 months after treatment.

### Oral administration of splicing compound using a formulated diet

The newly hybrid transgenic mice carrying the human *ELP1* transgene with FD mutation *TgFD9*;*Pax6cre;Elp1^flox/flox^*were fed a special diet containing 0.0008% of PTC-409680 (Purina Prolab 3000 with PTC-409680, #21032505A2i) corresponding to a daily consumption of 1.2 mg/kg/day.

### Retinal histology following ocular gene therapy with AAV2-U1a-*mElp1*

Retina flat mount dissection and histology were performed as previously described [39, 40]. In brief, Mice were euthanized with CO2 and the temporal surface of the eye was marked with a green tattoo dye (Ketchum Manufacturing) prior to enucleation. After the eyes were enucleated, a hole was made at the sclera-cornea margin and fixed overnight at 4°C in 4% PFA. Before antibody staining, the neural retinal layer was detached from the retinal pigment epithelium (RPE)/choroid layer. Rabbit anti-Brn3a (1:300; Synaptic Systems, Cat#411 003) was used as the primary antibody. Cy3 conjugated anti-rabbit (1:500, donkey) secondary antibody was purchased from Jackson Immuno Research. All antibodies were diluted in 1X PBS with 0.3% Triton X-100 and 5% bovine serum albumin (BSA, Cell Signaling Technology).

### Retinal histology following ocular gene therapy with AAV2-U1a-h*ELP1*

Mice were euthanized as stated above. Eyes were fixed at room temperature (RT) for 1 hour in 4% PFA. Retinae were then removed, with the temporal region marked by a small cut, and incubated in 4% PFA for an additional 15 minutes at RT. Tissue was permeabilized in 1X PBS containing 0.5% Triton X-100 for 30 minutes at RT. Non-specific binding was blocked by incubation with 5% BSA containing 0.5% Triton X-100 for 3 hours at RT. All retinae were incubated for 2 nights at 4°C in a primary antibody solution containing 2% BSA, 0.2% Triton X-100, an anti-Brn3 antibody (1:200, Santa Cruz, #sc-6026), and an anti-ELP1(IKAP) antibody (2µm/ml, Invitrogen, #PA5-111296, RRID AB_2856706). After overnight incubation at 4°C with secondary antibodies (Donkey anti-Goat IgG (H+L) Cross-Adsorbed Secondary Antibody, Alexa Fluor™ 647 and Donkey anti-Rabbit IgG (H+L) Highly Cross-Adsorbed Secondary Antibody, Alexa Fluor™ 568 from Thermo Fisher Scientific, catalog # A-21447, RRID AB_2535864 and catalog # A10042, RRID AB_2534017), four identical cuts were made to flatten the tissue into four lobes, and the tissue was mounted with Prolong Gold (Invitrogen).

### RGC quantification

Flat mounts were imaged and tiled using a Leica DM6 Thunder microscope at a 10X magnification. For the AAV2-h*ELP1* injected retinae, flat mount images were obtained with a Leica DMi8 THUNDER imaging system using the navigator spiral scan at a 10X magnification. IMARIS software (Oxford Instruments) was used to quantify the number of Brn3a+ cells. To identify Brn3a+ cells, the spots module was used. The detected diameter was set at 8 µm in the peripheral region determined by a 1 mm^2^ box placed at the edge of each quadrant in regions with the most cell death. If a specific region was damaged, the RGCs directly adjacent were counted. If any complication with the intravitreal injection, retinal harvest, or histology resulted in overall damaged retinal tissue, the sample was excluded and omitted from the analysis. The experimenters were blind to genotype and treatment group for all RGC quantification. The total peripheral cell count is the sum of the cells counted in the four peripheral regions. We analyzed 100% of retinae from each mouse but only report data from the peripheral retina because the *Pax6-Cre* expression is not expressed in the central retina. Therefore, the loss of Elp1 is restricted to the periphery [21].

### Statistics

All data are expressed as mean ± SEM. Column data are plotted as a scattered dot-bar plot to show variability, and n shows the sample size for each data set. GraphPad Prism (version 9) was used to make all graphs. P < 0.05 was considered to represent a significant difference. Following confirmation of normal distribution, data were analyzed using an unpaired two-tailed student’s t-test and/or one-way ANOVA followed by Tukey’s post hoc test for multiple comparisons.

For AAV2-h*ELP1* injection data, we visually checked normality, found a non-normal distribution, and confirmed with a Shapiro test (p = 7.89e^-8^). Due to non-normal distribution, we ran a Kruskal Wallis non-parametric test (p=0.0005). Followed by a post hoc Dunn test with Bonferroni correction, we found significance only between controls and *Pax6cre;Elp1^loxp/loxp^*mice, regardless of treatment. Since we had a small sample size with unequal variance and were most interested in the relationship between the uninjected *Pax6cre;Elp1^loxp/loxp^*and the *Pax6cre;Elp1^loxp/loxp^* receiving the highest viral titer (5.4×10^8^ vg/eye) we tested for normality and found a non-normal distribution. Therefore, using a Mann-Whitney U test we reject the null hypothesis that the distributions are equal and conclude there is a difference between uninjected *Pax6cre;Elp1^loxp/loxp^*and the *Pax6cre;Elp1^loxp/loxp^* that received the highest viral titer (p = 0.02).

All statistics were performed using GraphPad Prism (version 9) and R version 4.2.1 (R Core Team, 2022).

## Materials availability

The newly designed AAV2 vectors are publicly available from Vector Core in the University of Massachusetts Medical School Gene Therapy Center.

## Data availability

All data needed to assess the conclusions from the study are included in the paper and/or supplemental files. When applicable, individual data points are shown on each graph. Please address any requests for raw data counts to the corresponding author.

## Acknowledgments

We thank Dr. James Fox in the Animal Resource Center at Montana State University for his support and help with animal husbandry. We also thank Jamie Morgan and Luke Domanico at Montana State University for assisting with the cell culture and viral transduction experiments and Dr. Marc Mergy for surgical assistance.

## Funding

This work was supported by NIH NEI R21EY031130 and a grant from the Familial Dysautonomia Foundation.

## Additional Information

The authors declare competing financial interests.

Xin Zhao, Jana Narasimhan, and Marla Weetall are employees of PTC Therapeutics, Inc., a biotechnology company. In connection with such employment, the authors receive salary, benefits, and stock-based compensation, including stock options, restricted stock, other stock-related grants, and the right to purchase discounted stock through PTC’s employee stock purchase plan.

Susan A. Slaugenhaupt is a paid consultant to PTC Therapeutics and is an inventor on several U.S. and foreign patents and patent applications assigned to the Massachusetts General Hospital, including U.S Patents 8,729,025 and 9,265,766, both entitled “Methods for altering mRNA splicing and treating familial dysautonomia by administering benzyladenine,” filed on August 31, 2012 and May 19, 2014 and related to use of kinetin; and U.S. Patent 10,675,475 entitled, “Compounds for improving mRNA splicing” filed on July 14, 2017 and related to use of BPN-15477.

Elisabetta Morini and Susan A. Slaugenhaupt are inventors on an International Patent Application Number PCT/US2021/012103, assigned to Massachusetts General Hospital and PTC Therapeutics entitled "RNA Splicing Modulation" related to use of BPN-15477 in modulating splicing.

All other authors declare no competing interests.

## Author Information and Contributions

**Department of Microbiology and Cell Biology, Montana State University, Bozeman, MT**

Anastasia Schultz, Stephanann Costello, Heini Miettinen, Marta Chaverra, Colin King, Lynn George, and Frances Lefcort

**Department of Ophthalmology, Neurobiology & Gene Therapy Center, University of Massachusetts Chan Medical School, Worcester, MA**

Shun-Yun Cheng and Claudio Punzo

**Center for Genomic Medicine, Massachusetts General Hospital Research Institute, Boston, MA**

Emily Kirchner, Susan Slaugenhaupt, and Elisabetta Morini

**Department of Neurology, Massachusetts General Hospital Research Institute and Harvard Medical School, Boston, MA**

Susan Slaugenhaupt, and Elisabetta Morini

**Department of Biological and Physical Science, Montana State University Billings, Billings, MT**

Lynn George

**PTC Therapeutics, Inc., South Plainfield, NJ 07080**

Xin Zhao, Jana Narasimhan, and Marla Weetall

## Contributions

F.L., C.P., E.M., and S.S. supervised and oversaw the experimental approach and conceptualization. A.S., F.L., C.P. and E.M. designed this study. A.S., S.C., C.K., and H.M. conducted experiments. A.S., S.-Y.C., E.K., H.M., and M.C. collected the data. A.S. and S.-Y.C. performed data analysis. X.Z., J.N., and M.W. designed and contributed the chow containing the small splicing modulator. L.G. supervised animal husbandry and breeding strategies. A.S. and F.L. drafted the manuscript. A.S., S.-Y.C., E.K., H.M., M.C., L.G., S.C., C.P., F.L., and E.M. edited the manuscript. All authors read and approved the final manuscript.

