## Supplemental Figure 1 for "Reduction of retinal ganglion cell death in mouse models of familial dysautonomia using AAV-mediated gene therapy and splicing modulators"

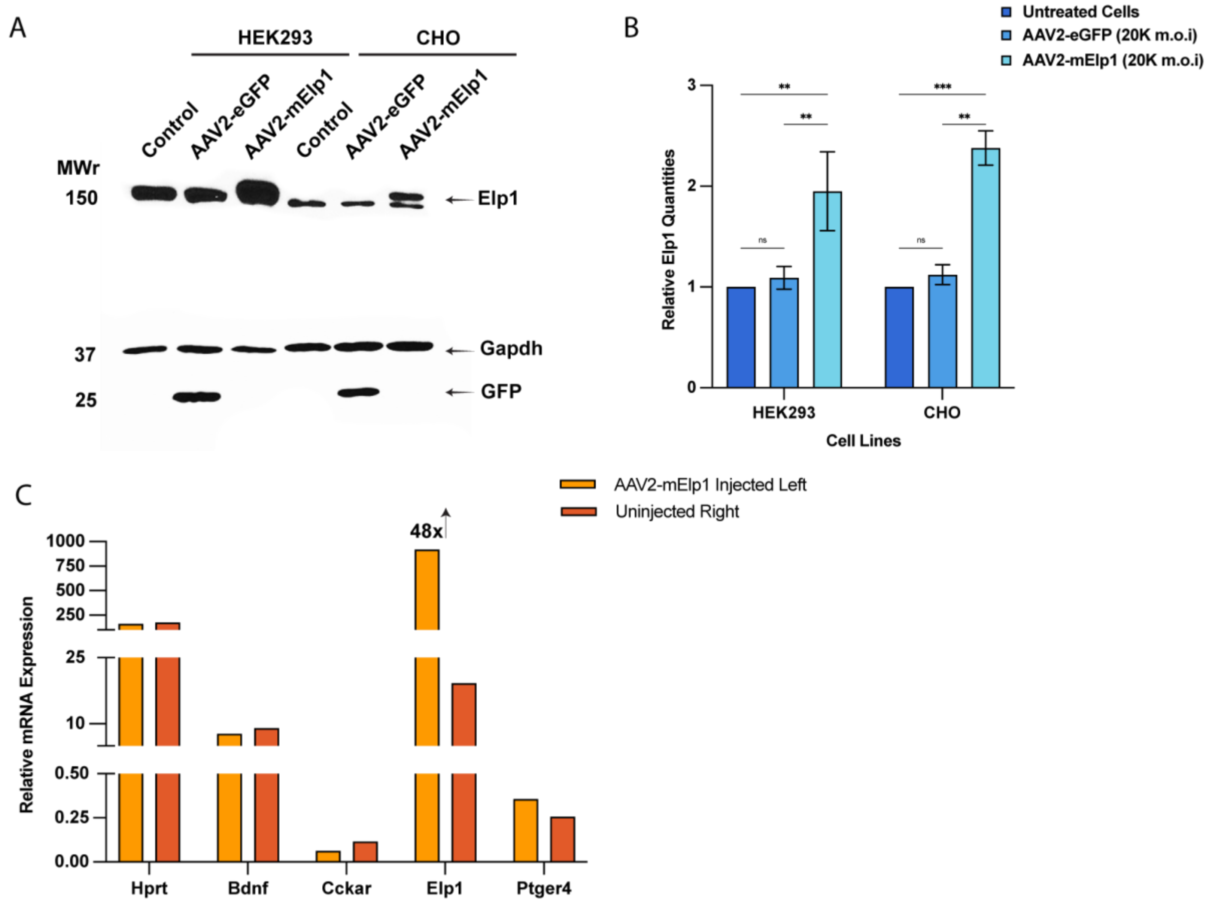


***Supplemental Figure 1:*** (A) Representative immunoblot showing the expression of murine Elp1 in two different cell lines 4 days post-transduction with AAV2-U1a-m*Elp1* and AAV2-U1a-eGFP. There is an increase in Elp1 expression in the transduced cells. 20µg of protein was loaded in each well, and GAPDH was used as the loading control. (B) Densitometric analysis of Elp1 protein in HEK293 and CHO cells transduced with AAV2-U1a-m*Elp1* compared to untreated cells (HEK293: **p = 0.004, 0.005; CHO: **p = 0.001, ***p = 0.0008, two-way ANOVA with Tukey's multiple comparisons follow-up test. The experiment was done in triplicate. (C) RT-qPCR analysis of *Elp1* expression in retinal homogenates 1 month after intravitreal injections of AAV2-U1a-m*Elp1*. Only Elp1 is strongly increased.
