## Supplemental Figure 2 for "Reduction of retinal ganglion cell death in mouse models of familial dysautonomia using AAV-mediated gene therapy and splicing modulators"

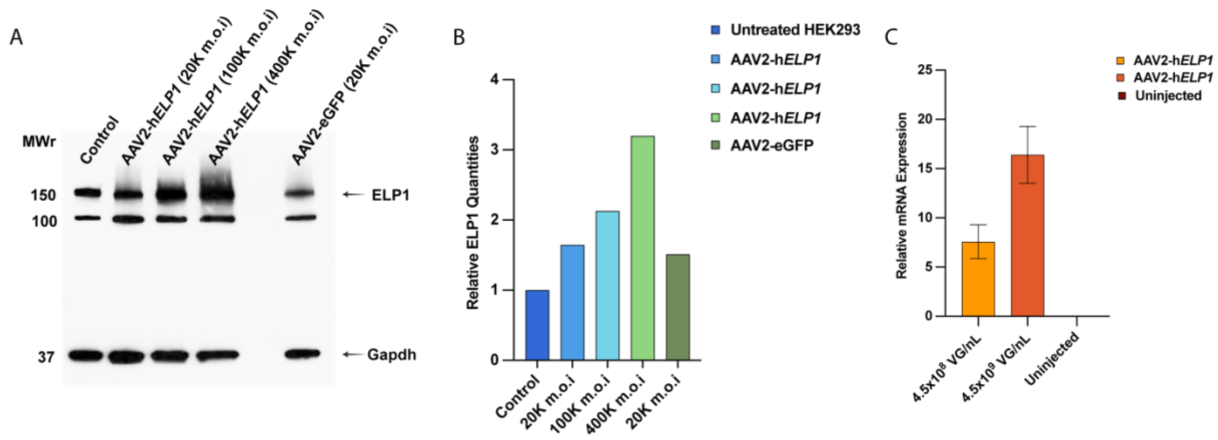


***Supplemental Figure 2:*** (A) Representative immunoblot showing the upregulation of human ELP1 in HEK293 cells 4 days post-transduction with AAV2-U1a-h*ELP1*. Note the gradual increase in ELP1 expression as the multiplicity of infection (m.o.i.) increases. Equal amounts of protein were loaded in each well, and GAPDH was used as the loading control. (B) Densitometric analysis of ELP1 protein in HEK293 cells transduced with AAV2-U1a-h*ELP1* compared to untreated cells. The experiment was done five times. (C) RT-qPCR analysis of *ELP1* expression in retinal homogenates (n=2) 1 month after intravitreal injections of AAV2-U1a-h*ELP1*. As expected, we saw no expression of the human ELP1 gene in the uninjected control homogenates, whereas the control mice receiving AAV2-U1a-h*ELP1* showed increasing levels of human ELP1 protein or mRNA.
