## Supplemental Table 1 for "Reduction of retinal ganglion cell death in mouse models of familial dysautonomia using AAV-mediated gene therapy and splicing modulators"

| Sequence Name | | Seq 5' to 3' | Accession No. |
|---|---|---|---|
| ELP1 | F | CTCTGCAGTCTCAGCACACA | NM_003640.5 |
| ELP1 | R | CTGCTCCAGGATTGGCTCAA | |
| Elp1 | F | GGTGACAGTCTTTCGGCAGA | NM_026079.3 |
| Elp1 | R | GATCAGCAGCCGGTAGGTAC | |
| Cckar | F | TGAACAAACGCTTTCGCCTG | NM_009827 |
| Cckar | R | TGGCTGTAGGAATACCGGGA | |
| Ptger4 | F | CACCACCTCGCTGAGAACTT | NM_008965 |
| Ptger4 | R | TCCTTTAGAGGCAGGCTCCT | |
| Hprt | F | TCAGTCAACGGGGGACATAAA | NM_013556 |
| Hprt | R | GGGGCTGTACTGCTTAACCAG | |
| Actb | F | AACCCTAAGGCCAACCGTGAA | NM_007393 |
| Actb | R | TCACGCACGATTTCCCTCTCA | |
| Bdnf | F | ACTGCAGTGGACATGTCTGG | NM_007540 |
| Bdnf | R | AGTTGGCCTTTGGATACCGG | |
